# Accumulation and retention of radioactive elements in biofilm communities surrounding the accident site of the Fukushima Daiichi Nuclear Power Plant

**DOI:** 10.1101/106211

**Authors:** Shigeharu Moriya, Hideaki Otsu, Kumiko Kihara, Yukari Kato, Misao Itouga, Kenji Sakata, Jun Kikuchi, Hiroshi Yamakawa, Jiro Suga

## Abstract

After the Fukushima Daiichi Nuclear Power Plant accident, various surveys have been performed to measure the extent of radioactive contamination in marine sediments, surface waters, plankton, and fish. However, the radioactive contamination of one of the most important ecological niches, biofilms, has not been investigated. Therefore, in this study, we sampled biofilms from sea floor stones around Hisanohama Port, which is less than 30 km south of the accident site, and then analyzed the microbial community structure and element profiles, including those of radioactive elements, of these biofilms in order to determine the accumulation and retention of radioactive elements in them. Our results showed that the biofilm samples contained relatively high levels of radioactive cesium even when the sampling was performed 8–11 months after the accident. Our results also suggested that the structure of the biofilm organismal community is related to the element profile of radioactive cesium. Thus, our study suggests that biofilms are a possible radioactive compound accumulator in the natural environment and that they can retain radioactive material for at least 8–11 months.

## Introduction

Following the Great East Japan Earthquake and catastrophic tsunami in March of 2011, the accident at the Fukushima Daiichi Nuclear Power Plant (FDNPP) led to the release of large amounts of radioactive materials into the atmosphere and marine environment. This release of radioactive elements from the FDNPP introduced a variety of radioisotopes, especially radiocesium (^134^Cs and ^137^Cs), into marine waters. Public monitoring showed that during several months after the first explosion at the FDNPP, a rapid decrease in radioisotopes was observed in the nearby seawater. A joint official survey team from the Tokyo Power Electric Company (TEPCO), Fukushima prefectural government, and the Japanese Ministry of Education, Culture, Sports, Science and Technology (MEXT) reported that the ^137^Cs contamination level in the seawater quickly decreased from over 10 k Bq/L to less than 100 Bq/L within a 2-month period. However, ^137^Cs in the ocean sediments was continuously observed over this period at levels between 100 Bq/dry kg and 1,000 Bq/dry kg at least until September 2012 (Kanda, 2012). This disproportional distribution of radioisotopes was also observed in marine fish by Wada et al., who reported that the retention time of radiocesium appears to be remarkably longer in the larvae of demersal fish such as Sebastes cheni than in surface fish such as Eugraulis japonica (Wada et al., 2013). These disproportional distributions of radioactivity suggest that the sea floor eco-system, including sedimental microbes and biofilms, may play role in the accumulation of radioactive elements.

Therefore, in this study, we analyzed the biofilm community on the surface of sea floor rocks with a focus on the element profiles, including those of radioactive elements. Our results showed that all the biofilm community contained and retained relatively high levels of radioactive cesium even when the sampling was performed 8–11 months after the accident. Our findings indicate that biofilm organism community structures are likely related to their element profiles, including that of radiocesium.

## Materials and Methods

### Sampling

Sampling was performed from November 11, 2011, to February 4, 2012. The sampling site was the ocean along the coastline of Hisanohama, Iwaki City, Fukushima, Japan. We collected samples 10 times within the established time period on November 11 (week 1), November 20 (week 2), November 26 (week 3), December 3 (week 4), December 10 (week 5), December 17 (week 6), January 14 (week 7), January 21 (week 8), January 29 (week 9), and February 4 (week 10). Sampling was performed at six different locations near Hisanohama Port: Area 1 (37.153762N 141.006167E, weeks 1 and 5), Area 2 (approximately 37.149863N 141.001017E, weeks 2, 8, and 9), Area 3 (approximately 37.146579N 141.002390E, week 4), Area 3′ (approximately 37.147503N 141.008570E, week 10), Area 4 (approximately 37.133887N 141.002390E, weeks 3 and 7), and Area 4′ (approximately 37.142953N 141.003592E, week 6); all of these sampling sites are located 30 km south of the Fukushima Nuclear Power Plant #1. Sampling was performed by divers using a self-contained underwater breathing apparatus (SCUBA) at a depth of 5–10 m. The divers sampled the bottom water and surface water (5 L) and collected bottom stones to obtain the biofilms. The samplings were performed with permission from Fukushima Prefecture and the fishery association of Iwaki City.

### Sample preparation

The seawater samples were immediately filtered using two 24-mm-wide 0.22-μm Durapore filters (Millipore, MA, USA) until they clogged. All the samples were then frozen at -20°C in a portable freezer prior to transportation back to the laboratory.

In the laboratory, the water samples were dried by boiling. The remaining salts (seawater salt) were allowed to settle into a U8 container, and the net weight (weight of container with sample – weight of empty container) and height of the samples were measured in the container.

In the case of biofilms, the bottom stones were rinsed with purified water and then brushed down with a clean brush using purified water. The obtained biofilm suspension was then divided into two aliquots. One aliquot was filtered through 24-mm-wide 0.22-μm Durapore filters (Millipore, MA, USA) and stored at -20°C for ribosomal RNA gene analysis, and the other one was dried at 105°C in an oven and then allowed to settle into a U8 container; the weight and height were then measured in the container for measurement of radioactivity.

### Measurement of radioactivity

The gamma-ray activity of the samples in the containers was measured using a Germanium (Ge) detector. The detector setup was identical to that described previously (HABA et al., 2012). The energy resolution of 2 keV full width at half maximum (FWHM) was achieved at 1332.5 keV gamma rays. The sample location and detector were surrounded with 15-cm lead blocks to reduce the background radiation during gamma ray spectroscopy. The detection efficiency of the gamma rays was calibrated with an accuracy of 2%–10% using a multiple gamma ray standard source, which ranged from 88–1836 keV. A calibrated ^134g^Cs source was also used to correct for the coincidence summing for radioactivity determinations of ^134g^Cs.

Every sample was contained in a U8 plastic container. The U8 container is a standard container in Japan that is used for measurements of absolute radioactivity with high accuracy using a Ge detector. The shape of the container is cylindrical, with a diameter of 47 mm and height of 60 mm. The volume is approximately 100 mL. The efficiency calibration source is available within the container with a uniform density.

Every sample was measured using the Ge detector for at least 8 h to obtain sufficient statistics for each gamma ray peak. The relative geometry between the sample and the Ge detector was carefully reproduced by a guide apparatus at the sample location.

Gamma rays from ^134^Cs and ^137^Cs were observed for the biofilm samples. The radioactivity levels for every isotope were determined from the yield of the gamma rays corrected using the detection efficiency of the Ge detector and a geometrical acceptance between the sample and the detector. The geometric difference in the height of the sample from the calibration source was simulated using several electromagnetic simulation codes, including Geant4 (Agostinelli et al., 2003; Allison et al., 2006) and EGS5 (Hirayama et al., 2005). The radioactivity of the samples were normalized by weight.

### Element analysis

To quantitatively analyze the content of 37 elements (^7^Li, ^9^Be, ^23^Na, ^24^Mg, ^27^Al, ^39^K, ^43^Ca, ^45^Sc, ^47^Ti, ^51^V, ^52^Cr, ^55^Mn, ^57^Fe, ^59^Co, ^60^Ni, ^63^Cu, ^66^Zn, ^85^Rb, ^88^Sr, ^98^Mo, ^111^Cd, ^133^Cs, ^139^La, ^140^Ce, ^141^Pr, ^146^Nd, ^152^Sm, ^153^Eu, ^158^Gd, ^159^Tb, ^164^Dy, ^165^Ho, ^166^Er, ^169^Tm, ^172^Yb, ^175^Lu, and ^202^Hg) in the biofilm samples for measurement of radioactivity, the samples were predigested with 5 mL of concentrated HNO3 for 1 h at room temperature. Next, the organic components were completely decomposed by wet-ashing using a microwave sample preparation system (Multi-Wave-3000, Perkin Elmer, MA USA) (Itouga et al., 2014). The digested samples were then brought up to a volume of 50 mL using MilliQ water (MQW) and filtered through 5B filter paper (Advantec, Tokyo, Japan). The concentrations of the mineral elements were determined by Inductively Coupled Plasma Mass Spectrometry (ICP-MS, NexION300, Perkin Elmer). For the ICP-MS analysis, a portion of the filtrated samples was diluted appropriately with 0.01 mol/L HCl (Itouga et al., 2014). The mineral nutrient concentration in the samples was calculated as a unit (mg/g in dry weight).

The obtained ICP-MS data were then normalized using the root-sum-of-squares (RSS) levels for each element. Count data of radioactive cesium (134Cs and 137Cs) were treated with the same calculation for normalization and then merged with the normalized ICP-MS dataset (element table).

### ssrDNA analysis

Genomic DNA was prepared from the stored biofilm samples by capturing on a 0.22-μm Durapore filter, followed by bead-beating with phenol:chloroform:isoamyl alcohol (PCI) extraction. Bead-beating was performed at 3000 rpm for 5 min (Micro Smash MS-100R, TOMY, Tokyo, Japan), and nucleic acids were then recovered by ethanol precipitation. PCR amplification was performed with the extracted genomic DNA using two primer sets for 16S ssrDNA, 515F/806R (Caporaso et al., 2012; Caporaso et al., 2011) and 18S ssrDNA, TAReuk454FWD1/TAReukREV3 (Stoeck et al., 2010). The primers were modified for Illumina sequencing with a multi-index kit for the Nextera XT sequencing library construction kit (Illumina, CA, USA). We also added a "read1" sequence (5′-TCG GCA GCG TCA GAT GTG TAT AAG AGA CAG -3′) to the forward primers and a "read2" sequence (5′-GTC TCG TGG GCT CGG AGA TGT GTA TAA GAG ACA G -3′) to the reverse primers. PCR was performed with a standard reaction of ExTaq DNA polymerase according to the manufacturer’s instructions (Takara Bio, Kyoto Japan) with 25 pmol of each primer/50 μL reaction mixture. The reactions were performed in a two-step PCR (10 cycles of 94°C for 30 s, 55°C for 45 s, and 72°C for 60 s, followed by 20 cycles of 94°C for 30 s and 72°C for 60 s) for 16S and a 3-step PCR (30 cycles of 94°C for 30 s, 55°C for 45 s and 72°C for 60 s) for 18S. The amplified products were then purified via 1% agarose gel electrophoresis. The obtained amplified products were applied to the NexteraXT (Illumina CA USA) sequencing library construction process without the “tagmentation” process.

A multi-index sequencing library was applied to the MiSeq sequencing kit v1 and read using Illumina MiSeq according to the instructions in the paired-end sequencing mode. Data were treated with MiSeq reporter software, and the reads were obtained from individual samples. After the extraction of the reads, the reads from weeks 4, 5, and 8 for the 18S ssrDNA were found to be insufficient for analysis. Therefore, subsequent analyses were performed using the reads from weeks 1, 2, 3, 6, 7, 9, and 10.

The obtained read data were then mapped onto the ribosomal RNA sequence library SILVA release 108 with the program package QIIME (Caporaso et al., 2010) using “Prefix-suffix operational taxonomy unit (OTU) picking,” and the absolute abundance OTU table at the family level was constructed. Count zero data were manually removed from the table, and the relative abundance values were calculated. The 16S and 18S data were treated independently and then merged into a single OTU table with the relative abundance values. Sequence data have been deposited in the DDBJ sequence read archive under the accession number DRA004367.

### Statistical analysis

For the statistical analysis, we used the program package R (R Core Team, 2014). First, we drew a heatmap chart with hierarchical cluster analysis (HCA). The heatmap.3 function in the GMD package, which is a package for non-parametric distance measurements between two discrete frequency distributions (Zhao et al., 2011), was used to draw the heatmap with HCA. Independent heatmaps were drawn from each element table and the OTU table with a two-axis HCA. A scaling option was used for the OTU table treatment according to the row direction but not for the elements table treatment.

Principal components analysis (PCA) was performed with the merged dataset, including both the OTU table and elements table, using the prcomp function in R. Score plots were drawn with principal component 1 (PC1; x-axis) and PC2 (y-axis). The loading plots for quadrants II and III were drawn manually.

## Results and Discussion

The gamma-ray energy spectra revealed significant peaks corresponding to natural radioisotope signals, such as that of ^40^K (data not shown), for all samples. Furthermore, we found significant peaks corresponding to ^134^Cs and ^137^Cs from the biofilm samples. However, we did not detect signals for ^134^Cs and ^137^Cs from the sea water salts and particles captured on filter (Table 1).

**Table 1.** Peak detection from samples.

| Material | Peak detection |  |
| --- | --- | --- |
| | $^{134}\text{Cs}$ | $^{137}\text{Cs}$ |
| Surface sea water salt | – | – |
| Bottom sea water salt | – | – |
| Captured particles in surface water | – | – |
| Captured particles in bottom water | – | – |
| Biofilms on bottom rocks | + | + |

It is possible that the ^134^Cs (half-life: 2.0652 years) and ^137^Cs (half-life: 30.1 years) released owing to prior nuclear weapons tests in the Pacific Ocean and the Chernobyl disaster have remained in the seabed around Japan. Therefore, we also analyzed samples collected from near Nishi-kawana in the Tokyo Bay (which is approximately 280 kilometers from the FDNPP) as controls, but found no significant signals for the radioisotopes ^134^Cs and ^137^Cs from these control samples. Therefore, we concluded that the radioisotopes detected in the samples around Hisanohama port in this study originated from the FDNPP disaster rather than from the environmental background or from past nuclear weapons tests. These results indicated that the biofilms on the sea floor around Hisanohama retained radioactive material for at least 8–11 months after the FDNPP accident.

However, the observed radioactivity level was different among the different biofilm samples (Table 2). Therefore, we analyzed the relationship between the microbial consortia within the biofilms and the elemental composition of the biofilms. To perform this analysis, quantified element data were combined with radioactivity data for radiocesium (^134^Cs and ^137^Cs) and utilized for statistical analysis. Ribosomal RNA genes were sequenced and used to construct a dataset for the statistical analysis.

**Table 2.**
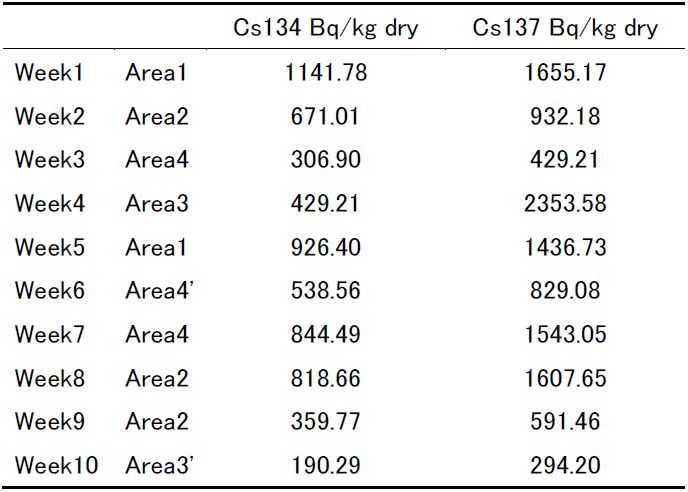
Radioactivity of biofilm samples.

|  |  | Cs134 Bq/kg dry | Cs137 Bq/kg dry |
| --- | --- | --- | --- |
| Week1 | Area1 | 1141.78 | 1655.17 |
| Week2 | Area2 | 671.01 | 932.18 |
| Week3 | Area4 | 306.90 | 429.21 |
| Week4 | Area3 | 429.21 | 2353.58 |
| Week5 | Area1 | 926.40 | 1436.73 |
| Week6 | Area4' | 538.56 | 829.08 |
| Week7 | Area4 | 844.49 | 1543.05 |
| Week8 | Area2 | 818.66 | 1607.65 |
| Week9 | Area2 | 359.77 | 591.46 |
| Week10 | Area3' | 190.29 | 294.20 |

A HCA heat map chart from the dataset of elements and radiocesium (Fig. 1) showed relatively higher content of radiocesium, Cr, Ni, Sc, Rb, Li, and cesium (Cs cluster) for the week 1 sample than for other samples. Another remarkable profile from the elemental analysis was observed in the week 3 sample, where high relative amounts of lanthanide elements were detected (Lanth cluster).

**Figure 1.**
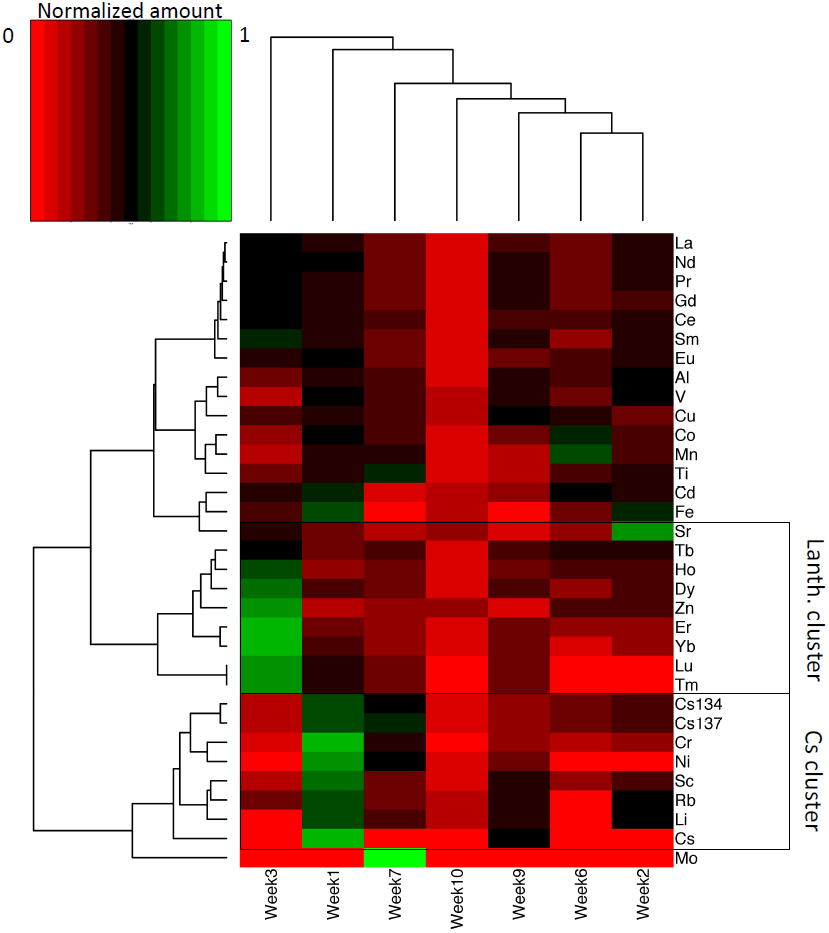
Heatmap chart of the elements and radioactivity data from the biofilm samples based on hierarchical clustering analysis (HCA). The heatmap chart was sorted by HCA for both axes (elements axis and sample axis). Boxes indicate clusters of radiocesium (Cs cluster) or lanthanides (Lanth cluster).

A PCA score plot (Fig. 2A) of the merged dataset between the elements/radiocesium and the microbial consortia showed that the week 1 sample was placed in the PC1 negative-PC2 positive direction, and the week 3 sample was placed in the PC1 negative-PC2 negative direction. As mentioned above, the week 1 sample contained high relative amounts of Cs cluster signals, and the week 3 sample contained high relative amounts of the Lanth cluster signals. Consistent with these findings, the PCA loading plot showed that the Cs cluster was placed in the PC1 negative-PC2 positive direction (Fig. 2B upper panel, Table S1), and the lanthanide cluster was placed in the PC1 negative-PC2 negative direction (Fig. 2B lower panel, Table S2). The results of the HCA heat map chart and PCA suggested that a "cesium-philic" biofilm occurred at the sampling point in the week 1 sample, and a "lanthanide-philic" biofilm occurred at the sampling point in the week 3 sample. The PCA results from the datasets of both the elements and the microbes showed candidate organisms in the "cesium-philic" and "lanthanide-philic" biofilms on the loading plots, because those plots were placed in same loading direction, e.g., PC1 negative-PC2 positive (Cs cluster included) or PC1 negative-PC2 negative (Lanth cluster included, Fig. 2C, Table S1 and S2). Indeed, the HCA heat map of the organismal consortia showed that the PC1 negative-PC2 negative–directed organisms mainly appeared in the week 1 column (Fig. 3 blue dots). The same tendency was observed with the Lanth cluster organisms (Fig. 3 green dots). These results suggested that these two OTU sets are positively correlated with the Lanth cluster (week 3 sample) and Cs cluster (week 1 sample) elements.

**Figure 2.**
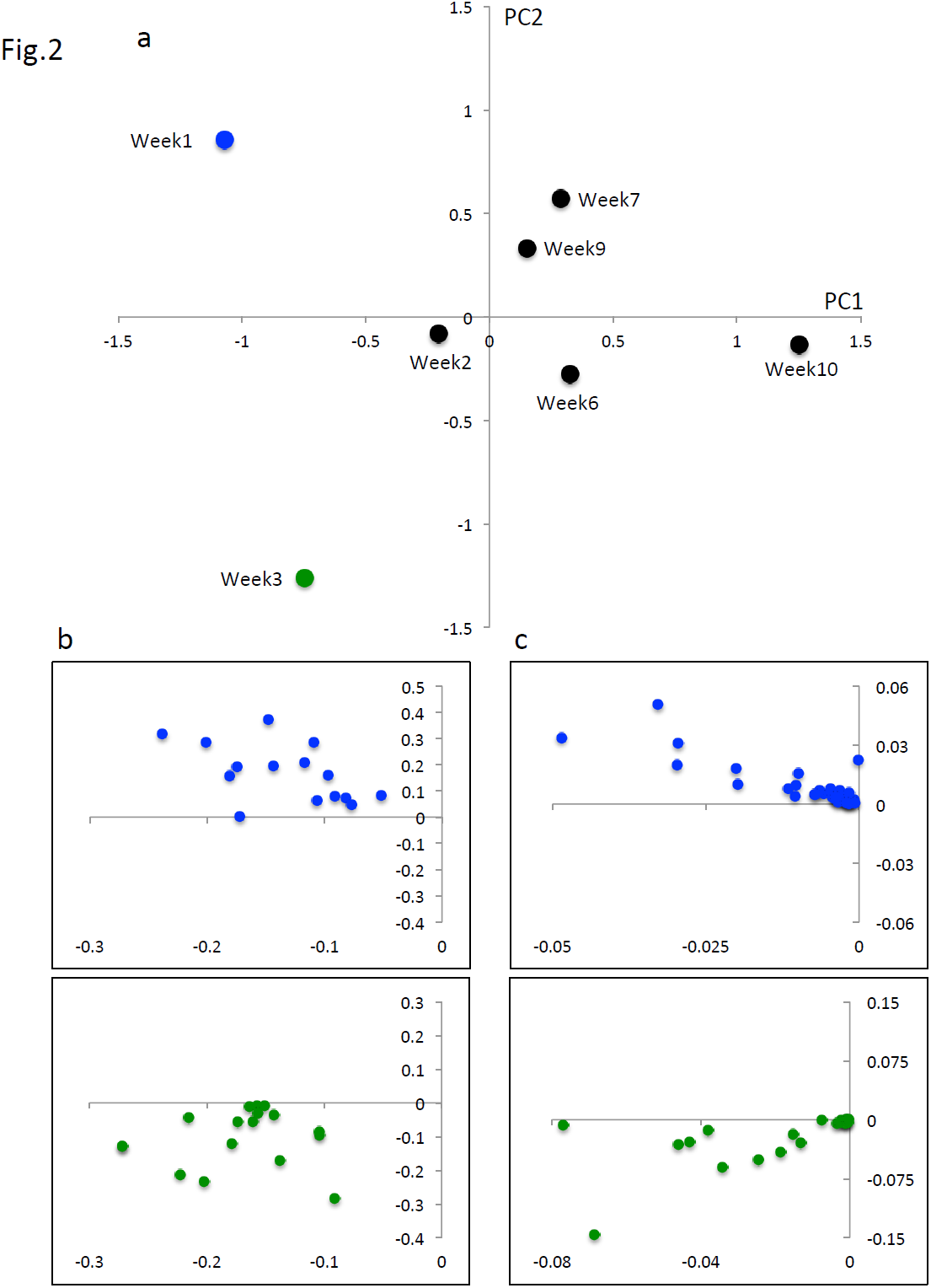
Principal component analysis (PCA) results for the sampling sites with microbial consortia data and element data with radiocesium. Panel A is the PCA score plot among sampling sites. Each dot indicates one sampling site. Panel B indicates the loading plot of the elements data. The upper panel shows the plot of elements that contribute to the PC1-negative and PC2-positive directions (supplementary table 1), indicating that they contribute to separate week 1 samples in the PC1-negative and PC2-positive directions. The lower panel shows the plot of elements that contribute to the PC1-negative and PC2-negative directions (supplementary table 2), indicating that they contribute to separate week 3 samples in the PC1-negative and PC2-negative directions in the PCA score plot. Panel C indicates the loading plot of the microbial consortia data. The upper panel displays the plot of operational taxonomic units (OTUs) that contribute to the PC1-negative and PC2-positive direction (supplementary table 1), indicating that they contribute to separate week 1 samples in the PCA score plot. The lower panel shows the plot of the OTUs that contribute to the PC1-negative and PC2-negative directions (supplementary table 2), indicating that they contribute to separate week 3 samples in the PCA score plot.

**Figure 3.**
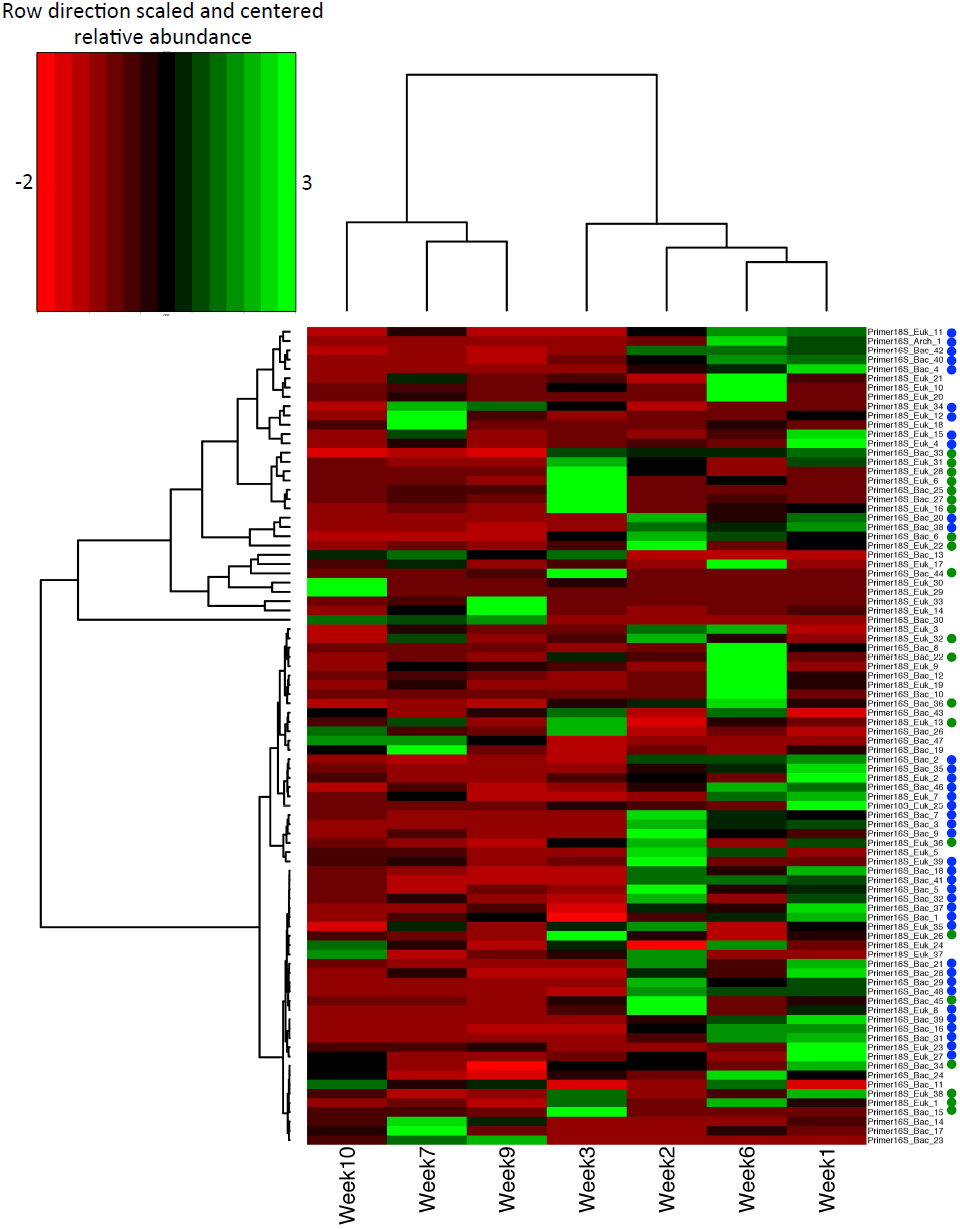
Heatmap chart of microbial consortia data for biofilms with HCA.The heatmap chart has been sorted by HCA for both axes (OTU axis and sample axis). Green and blue dots indicate contributing OTUs for the PC1-negative and PC2-positive directions and the PC1/PC2-negative direction, respectively. The relative intensity value has been scaled and centered in the row direction.

Table 3 lists the OTU set contents that correlated with Cs cluster elements, including radiocesium, and the relative intensity of each OTU in the biofilm community. The most abundant taxa in this OTU set were Hexacorallia and Oligochaete among the eukaryotes. For bacteria, high relative intensities of Bacteroidetes, Rhodobacteraceae, and Gammaproteobacteria were observed. Although a primer set that did not target Archaea was used, archeal sequences were obtained in this work by sequencing amplicons generated using the 16S ribosomal RNA primer set. An uncultured Crenarcheota in marine group I was observed to be one of the dominant OTUs with a correlation with Cs cluster elements. Previously, Alpha- and Gamma-proteobacteria have been obtained from biofilms on the surface of nuclear fuel pools (Sarró et al., 2005). According to Sarro et al. (Sarró et al., 2005), biofilms obtained from nuclear fuel pools accumulate ^60^Co. In our study, cobalt accumulation in the week 1 sample was suggested by the PCA data (Fig. 2B upper panel and Table S1) despite a lack of ^60^Co signal during gamma-ray spectroscopy. Microbial community retrieved from nuclear fuel pools have been described previously as capable of accumulating both ^60^Co and ^137^Cs (Tišáková et al., 2012). Our results suggest that the biofilm sampled in week 1 in our study presents similar characteristics to the microbes described by Sarro et al. and Tisakova et al. in the above-mentioned previous studies.

**Table 3.**
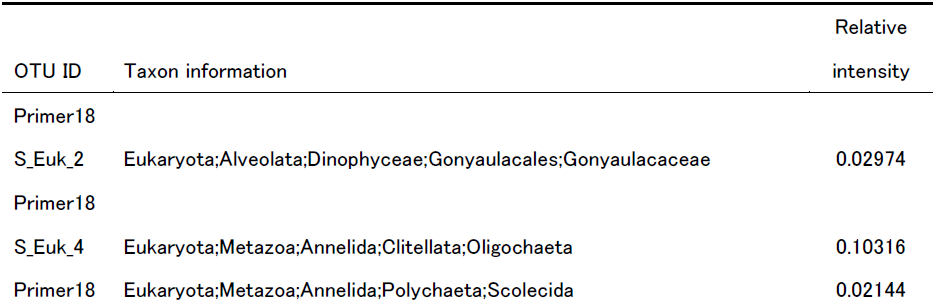
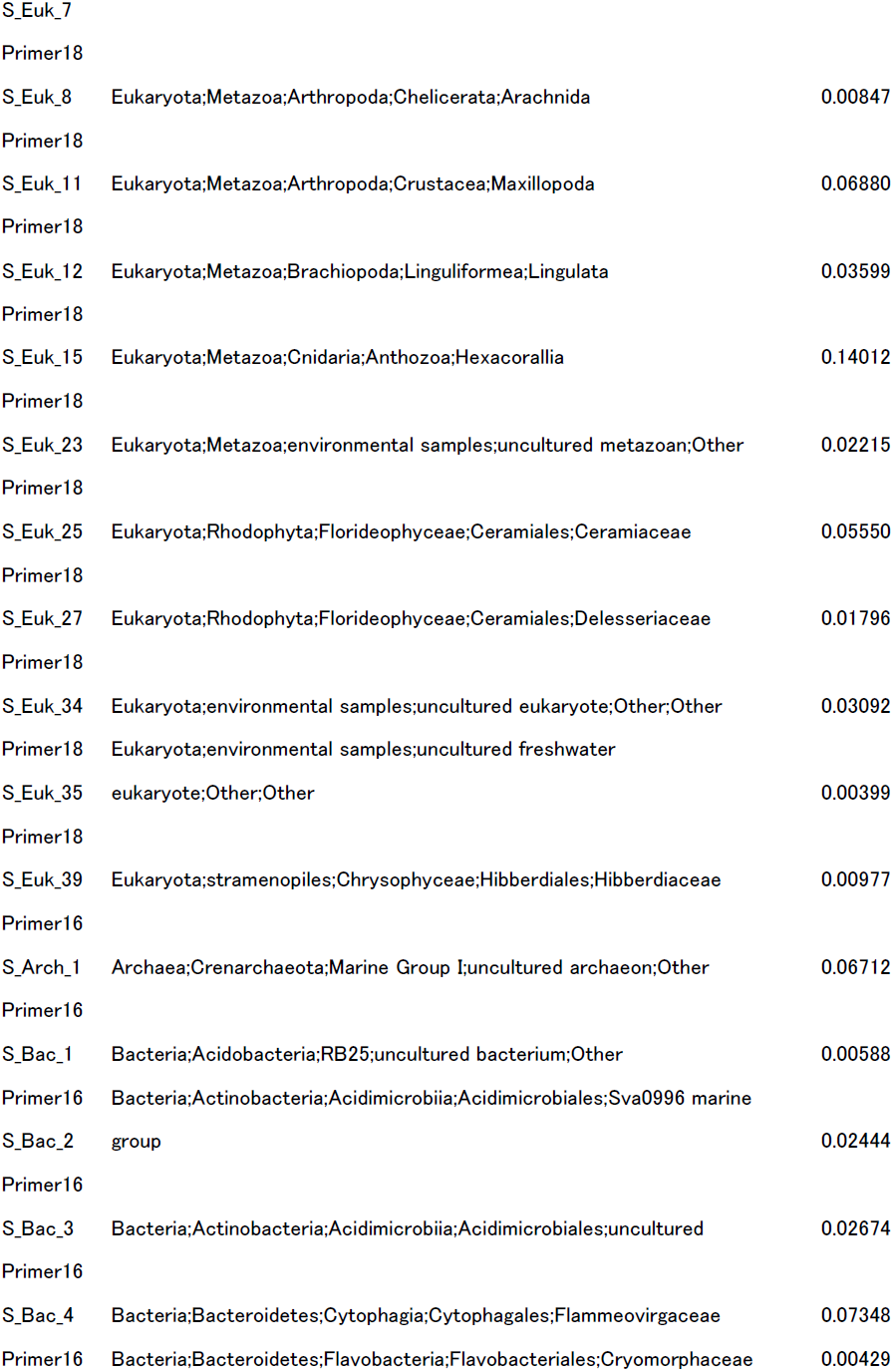
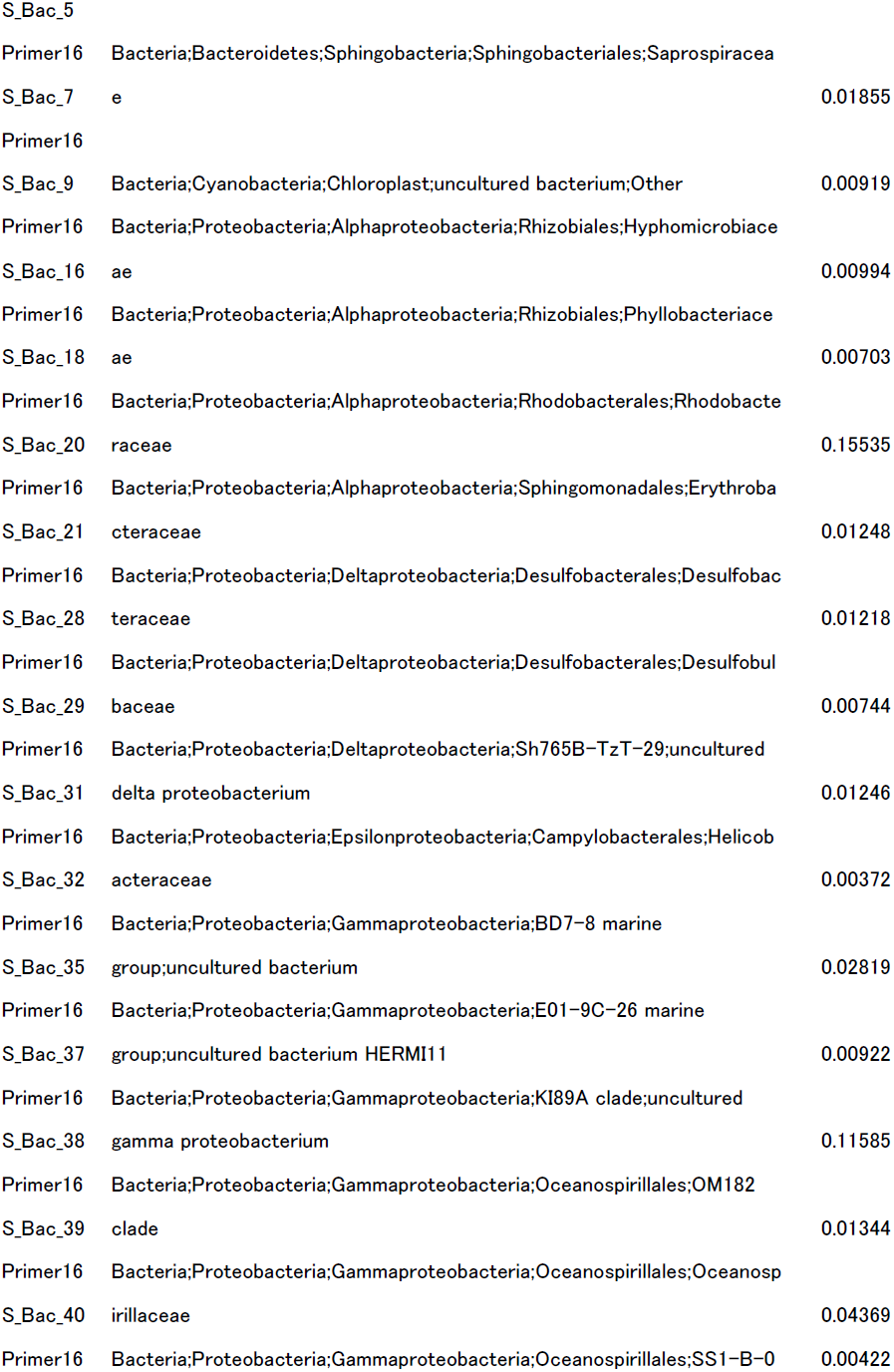
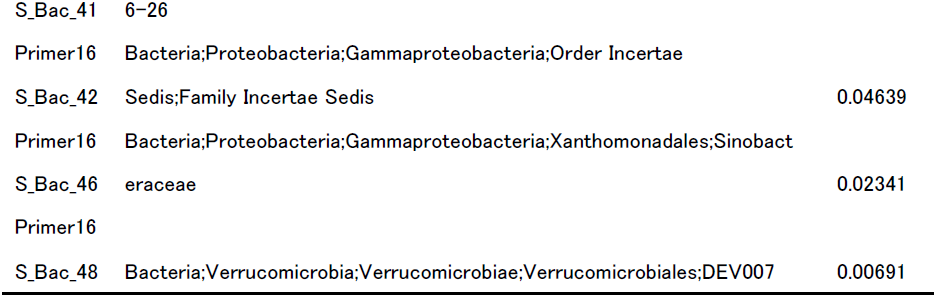
Cs cluster taxonin week 1 sample.

| OTU ID | Taxon information | Relative intensity |
| --- | --- | --- |
| Primer18 |  |  |
| S_Euk_2 | Eukaryota;Alveolata;Dinophyceae;Gonyaulacales;Gonyaulacaceae | 0.02974 |
| Primer18 |  |  |
| S_Euk_4 | Eukaryota;Metazoa;Annelida;Clitellata;Oligochaeta | 0.10316 |
| Primer18 | Eukaryota;Metazoa;Annelida;Polychaeta;Scolecida | 0.02144 |
| S_Euk_7 |  |  |
| Primer18 |  |  |
| S_Euk_8 | Eukaryota;Metazoa;Arthropoda;Chelicerata;Arachnida | 0.00847 |
| Primer18 |  |  |
| S_Euk_11 | Eukaryota;Metazoa;Arthropoda;Crustacea;Maxillopoda | 0.06880 |
| Primer18 |  |  |
| S_Euk_12 | Eukaryota;Metazoa;Brachiopoda;Linguliformea;Lingulata | 0.03599 |
| Primer18 |  |  |
| S_Euk_15 | Eukaryota;Metazoa;Cnidaria;Anthozoa;Hexacorallia | 0.14012 |
| Primer18 |  |  |
| S_Euk_23 | Eukaryota;Metazoa;environmental samples;uncultured metazoan;Other | 0.02215 |
| Primer18 |  |  |
| S_Euk_25 | Eukaryota;Rhodophyta;Florideophyceae;Ceramiales;Ceramiaceae | 0.05550 |
| Primer18 |  |  |
| S_Euk_27 | Eukaryota;Rhodophyta;Florideophyceae;Ceramiales;Delesseriaceae | 0.01796 |
| Primer18 |  |  |
| S_Euk_34 | Eukaryota;environmental samples;uncultured eukaryote;Other;Other | 0.03092 |
| Primer18 | Eukaryota;environmental samples;uncultured freshwater |  |
| S_Euk_35 | eukaryote;Other;Other | 0.00399 |
| Primer18 |  |  |
| S_Euk_39 | Eukaryota;stramenopiles;Chrysophyceae;Hibberdiales;Hibberdiaceae | 0.00977 |
| Primer16 |  |  |
| S_Arch_1 | Archaea;Crenarchaeota;Marine Group I;uncultured archaeon;Other | 0.06712 |
| Primer16 |  |  |
| S_Bac_1 | Bacteria;Acidobacteria;RB25;uncultured bacterium;Other | 0.00588 |
| Primer16 | Bacteria;Actinobacteria;Acidimicrobiia;Acidimicrobiales;Sva0996 marine |  |
| S_Bac_2 | group | 0.02444 |
| Primer16 |  |  |
| S_Bac_3 | Bacteria;Actinobacteria;Acidimicrobiia;Acidimicrobiales;uncultured | 0.02674 |
| Primer16 |  |  |
| S_Bac_4 | Bacteria;Bacteroidetes;Cytophagia;Cytophagales;Flammeovirgaceae | 0.07348 |
| Primer16 | Bacteria;Bacteroidetes;Flavobacteria;Flavobacteriales;Cryomorphaceae | 0.00429 |
| S_Bac_5 |  |  |
| Primer16 | Bacteria;Bacteroidetes;Sphingobacteria;Sphingobacteriales;Saprospiracea |  |
| S_Bac_7 | e | 0.01855 |
| Primer16 |  |  |
| S_Bac_9 | Bacteria;Cyanobacteria;Chloroplast;uncultured bacterium;Other | 0.00919 |
| Primer16 | Bacteria;Proteobacteria;Alphaproteobacteria;Rhizobiales;Hyphomicrobiace |  |
| S_Bac_16 | ae | 0.00994 |
| Primer16 | Bacteria;Proteobacteria;Alphaproteobacteria;Rhizobiales;Phyllobacteriace |  |
| S_Bac_18 | ae | 0.00703 |
| Primer16 | Bacteria;Proteobacteria;Alphaproteobacteria;Rhodobacterales;Rhodobacte |  |
| S_Bac_20 | raceae | 0.15535 |
| Primer16 | Bacteria;Proteobacteria;Alphaproteobacteria;Sphingomonadales;Erythroba |  |
| S_Bac_21 | cteraceae | 0.01248 |
| Primer16 | Bacteria;Proteobacteria;Deltaproteobacteria;Desulfobacterales;Desulfobac |  |
| S_Bac_28 | teraceae | 0.01218 |
| Primer16 | Bacteria;Proteobacteria;Deltaproteobacteria;Desulfobacterales;Desulfobul |  |
| S_Bac_29 | baceae | 0.00744 |
| Primer16 | Bacteria;Proteobacteria;Deltaproteobacteria;Sh765B-TzT-29;uncultured |  |
| S_Bac_31 | delta proteobacterium | 0.01246 |
| Primer16 | Bacteria;Proteobacteria;Epsilonproteobacteria;Campylobacterales;Helicob |  |
| S_Bac_32 | acteraceae | 0.00372 |
| Primer16 | Bacteria;Proteobacteria;Gammaproteobacteria;BD7-8 marine |  |
| S_Bac_35 | group;uncultured bacterium | 0.02819 |
| Primer16 | Bacteria;Proteobacteria;Gammaproteobacteria;E01-9C-26 marine |  |
| S_Bac_37 | group;uncultured bacterium HERMI11 | 0.00922 |
| Primer16 | Bacteria;Proteobacteria;Gammaproteobacteria;KI89A clade;uncultured |  |
| S_Bac_38 | gamma proteobacterium | 0.11585 |
| Primer16 | Bacteria;Proteobacteria;Gammaproteobacteria;Oceanospirillales;OM182 |  |
| S_Bac_39 | clade | 0.01344 |
| Primer16 | Bacteria;Proteobacteria;Gammaproteobacteria;Oceanospirillales;Oceanosp |  |
| S_Bac_40 | irillaceae | 0.04369 |
| Primer16 | Bacteria;Proteobacteria;Gammaproteobacteria;Oceanospirillales;SS1-B-0 | 0.00422 |
| S_Bac_41 | 6-26 |  |
| Primer16 | Bacteria;Proteobacteria;Gammaproteobacteria;Order Incertae |  |
| S_Bac_42 | Sedis;Family Incertae Sedis | 0.04639 |
| Primer16 | Bacteria;Proteobacteria;Gammaproteobacteria;Xanthomonadales;Sinobact |  |
| S_Bac_46 | eraceae | 0.02341 |
| Primer16 |  |  |
| S_Bac_48 | Bacteria;Verrucomicrobia;Verrucomicrobiae;Verrucomicrobiales;DEV007 | 0.00691 |

## Conclusions

In this study, we found that biofilms retain a relatively high amount of radioactive cesium in their microbial communities even after more than half a year from the release of radioactive elements into oceans due to nuclear power plant accidents. Our statistical analysis also indicated that different community structures of biofilms show different affinities for radiocesium. Our results suggested biofilms as a point of entry for radioactive elements into the eco-system owing to long-term retention of radiocesium in the biofilm communities.In the future, further research on the accumulation and retention of radioactive elements by marine biofilm communities needs to be undertaken to obtain information that can be used for developing ways to eliminate radioactive contamination and for monitoring of environmental pollution by radiocesium.

## Acknowledgements

This work was supported by grants for the RIKEN Incentive Research Project (FY2011) “Emergency monitoring for possible cycling of radio active compounds via oceanic food web” awarded to SM and HO and a MEXT Grant-in-Aid for Scientific Research on Innovative Areas (No. 23117003) awarded to SM and KK. We also wish to thank the Iwaki City Fishery Association for permitting the sampling for our study and for providing the fishing vessel Shyo-ei-maru and supporting crew belonging to the abalone section of the Iwaki Fishery Association for our study. We are also grateful to Toshiyuki Suzuki for overseeing and ensuring the diving safety of SM. The authors also wish to acknowledge the profound academic comments and proofreading provided by R. Craig Everroad and Diogo M. O. Ogawa.

## Supplementary tables.xlsx

Supplementary table 1

Supplementary table 2

